# Enhanced Ba^2+^-sensitive inward rectifying potassium conductance reduces intrinsic excitability of layer 2/3 pyramidal neurons in the primary auditory cortex of Fmr1 knockout mice

**DOI:** 10.64898/2026.02.03.703439

**Authors:** Claudio Moreno, Denise Riquelme, Christian Cea-del-Rio, Elias Leiva-Salcedo

**Affiliations:** Department of Biology, Faculty of Chemistry and Biology, University of Santiago, Santiago, Chile; PhD program in Neuroscience, University of Santiago of Chile; Centro de Investigación Biomédica y Aplicada (CIBAP), Escuela de Medicina, Facultad de Ciencias Médicas, Universidad de Santiago de Chile, Santiago, Chile

## Abstract

Fragile X syndrome (FXS) is frequently associated with auditory hypersensitivity and altered cortical processing, yet the intrinsic ionic mechanisms shaping auditory cortex excitability remain incompletely defined. Here, we tested whether subthreshold conductances contribute to compensatory changes in intrinsic excitability in layer 2/3 (L2/3) pyramidal neurons of the primary auditory cortex (AC) in Fmr1 knockout (Fmr1-KO) mice. We performed wholecell patchclamp recordings in acute slices from juvenile mice (P28–P42) and quantified passive properties, firing output, synaptic potentials, and subthreshold currents under pharmacological isolation. Compared with wild type (WT), Fmr1-KO L2/3 neurons displayed a more hyperpolarized resting membrane potential, reduced input resistance, elevated rheobase, prolonged first-spike latency, and reduced firing across depolarizing steps, consistent with a hypoexcitable state. Bath application of BaCl_2_ (60 μM) depolarized the membrane, increased input resistance, and restored firing output and rheobase toward WT levels, indicating that a Ba^2+^-sensitive potassium conductance strongly constrains excitability in Fmr1-KO neurons. Voltage-clamp recordings revealed a larger Ba^2+^-sensitive inwardly rectifying (Kir-like) current in Fmr1-KO neurons, supporting increased functional Ba^2+^-sensitive inwardly rectifying conductance. In contrast, blocking I_h_ with ZD7288 (10 μM) produced modest changes in passive properties but induced genotype dependent effects on excitability and enhanced synaptic activity and EPSP summation preferentially in Fmr1-KO neurons, consistent with a role for I_h_ in input filtering rather than setting basal conductance. Together, these findings identify enhanced Ba^2+^-sensitive potassium conductance as a primary determinant of the Fmr1-KO subthreshold conductance state in AC L2/3 pyramidal neurons, suggesting an intrinsic homeostatic mechanism that stabilizes output in the presence of elevated excitatory drive.

## 1. INTRODUCTION

Fragile X syndrome (FXS) is the most common inherited cause of intellectual disability and autism (Hagerman et al., 2017; Dahlhaus, 2018). It arises from a CGG trinucleotide expansion (>200 repeats) in the promoter region of the Fmr1 gene, leading to hypermethylation and transcriptional silencing of Fragile X Messenger Ribonucleoprotein 1 (FMRP) (Fernández et al., 2013; Brager & Johnston, 2014; Ferron, 2016). FMRP is an RNA-binding protein that regulates the dendritic transport and local translation of specific mRNAs, thereby controlling the expression of many synaptic and ion channel proteins (Richter & Zhao, 2021).

Patients with FXS frequently exhibit auditory hypersensitivity, reflecting abnormal processing of acoustic stimuli (Rotschafer & Razak, 2014; Ethridge et al., 2017, 2019; Bowen et al., 2020; Razak et al., 2021; Holley et al., 2022). In mouse models, this phenotype has been linked to increased excitability in subcortical auditory centers, which in turn induces compensatory homeostatic adaptations in cortical circuits (Garcia-Pino et al., 2017; Booker et al., 2019). Yet, the specific ionic mechanisms underlying this cortical compensation remain poorly defined.

Following the loss of FMRP, neuronal excitability is profoundly affected due to disrupted regulation of ion channel expression and function (Contractor et al., 2015; Bülow et al., 2019; Deng & Klyachko, 2021). Among these, subthreshold currents (active below the action potential threshold) play essential roles in determining how neurons maintain or adjust their resting membrane potential and responsiveness to synaptic input. Two such currents, the inwardly rectifying potassium current (Kir) and the hyperpolarization-activated cation current (I_h_), exert opposing effects on membrane excitability.

Kir currents stabilize the resting membrane potential by promoting potassium influx at hyperpolarized voltages and decreasing input resistance, thereby filtering out weak synaptic inputs (Day et al., 2005a; Pignatelli et al., 2019). Enhanced Kir activity shifts neurons into a high-conductance, hypoexcitable state that requires stronger inputs to reach firing threshold (Day et al., 2005b). Conversely, I_h_ currents, generated by hyperpolarization-activated cyclic nucleotide-gated (HCN) channels, depolarize the membrane through Na^+^ influx and promote subthreshold oscillations in the resting membrane potential (Campanac et al., 2008; Pavlov et al., 2011), and in dendrites controlling the duration of the EPSP. In cortical pyramidal neurons, HCN channels are distributed along the apical dendrite, shaping signal integration and dendritic filtering properties (Lörincz et al., 2002).

FMRP is known to interact with HCN channel mRNAs, and its loss reduces dendritic trafficking of these transcripts, potentially enhancing dendritic excitability (Brager & Johnston, 2014; Contractor et al., 2015; Brandalise et al., 2020). However, while altered I_h_ has been implicated in some cortical regions, evidence from the auditory cortex remains inconclusive. In contrast, the contribution of Kir channels to FXS-related changes in cortical excitability has received little attention. Although FMRP regulates astrocytic Kir4.1 expression (Bataveljic et al., 2024), and pharmacological Kir blockade rescues long-term potentiation deficits in hippocampal neurons of Fmr1-KO mice (Ordemann et al., 2021), the role of neuronally expressed Kir channels and their contribution to cortical auditory circuits dysfunction in Fmr1-KO mice has not been established.

Here, we examined layer 2/3 (L2/3) pyramidal neurons in the primary auditory cortex (AC) of Fmr1-KO mice to determine how alterations in subthreshold currents contribute to their intrinsic excitability. We found that these neurons exhibit an increased Ba^2+^-sensitive inward rectifying potassium current, while I_h_ baseline contribution to resting conductance is not significantly altered in Fmr1-KO. This enhanced Ba^2+^-sensitive inward rectifying potassium current activity is associated with a more hyperpolarized resting membrane potential, lower input resistance, and reduced firing frequency, reflecting a homeostatic shift toward a hypoexcitable state. Pharmacological inhibition of Ba^2+^-sensitive potassium conductance restored excitability to wild-type levels, suggesting that increased Ba^2+^-sensitive potassium conductance functions as a compensatory mechanism to stabilize neuronal output in the face of excessive excitatory drive.

## 2. METHODS

### 2.1 Animals

All procedures were conducted in accordance with protocols approved by the Ethical Committee of the University of Santiago de Chile and adhered to the guidelines established by the National Agency of Research and Development (ANID). Male C57BL/6J and B6.129P2-Fmr1^tm1Cgr^/J mice were housed in a temperature and humidity-controlled facility under a 12 h light/dark cycle, with ad libitum access to food and water.

### 2.2 Electrophysiological Recordings

#### 2.2.1 Brain Slice Preparation

Whole-cell patch-clamp recordings were performed in layer 2/3 (L2/3) pyramidal neurons of the primary auditory cortex from wild-type (WT; C57BL/6J) and Fmr1-KO (B6.129P2-Fmr1^tm1Cgr^/J) mice, between 28 and 42 days old. Animals were deeply anesthetized with 4% isoflurane and transcardially perfused with ice-cold, oxygenated, high-magnesium artificial cerebrospinal fluid (high-Mg^2+^ ACSF) containing (in mM): 124 NaCl, 2.5 KCl, 5 MgCl_2_, 0.5 CaCl_2_, 1.25 NaH_2_PO_4_, 25 NaHCO_3_, and 11 glucose (pH 7.4; 300 ± 5 mOsm/kg), continuously bubbled with 95% O_2_ / 5% CO_2_.

Brains were rapidly extracted following a midline and lateral skull incision and immersed in the same ice-cold high-Mg^2+^ ACSF. Coronal slices (350 μm thick) containing the primary auditory cortex were cut using a vibrating microtome (VT1000S, Leica Microsystems). Slices were incubated for 30 min at 37 °C in oxygenated ACSF containing (in mM): 125 NaCl, 2.5 KCl, 1.3 MgCl_2_, 2.5 CaCl_2_, 1.25 NaH_2_PO_4_, 25 NaHCO_3_, and 11 glucose (pH 7.4; 300 ± 5 mOsm/kg), and then maintained at room temperature (22-24 °C) until use. For recordings, slices were transferred to a submerged recording chamber mounted on a Zeiss Axio Examiner A1 DIC microscope and continuously perfused with oxygenated ACSF (1-3 mL/min) at 34 ± 2 °C. Pyramidal neurons were visually identified based on soma morphology, apical dendrite projection, and regular-spiking firing patterns.

#### 2.2.2 Current Clamp Recordings

Borosilicate glass pipettes (3-6 MΩ; Sutter Instruments, USA) were filled with an intracellular solution containing (in mM): 120 K-gluconate, 10 KCl, 8 NaCl, 10 HEPES, 0.5 EGTA, 4 Mg-ATP, 0.3 Na-GTP, and 10 phosphocreatine (pH 7.2, adjusted with KOH; 295-305 mOsm/kg).

Intrinsic excitability was measured using somatic current injections from -250 to +500 pA in 50 pA increments (500 ms duration) delivered at 0.2 Hz. Each sweep included -50 and +50 pA square pulses to monitor series resistance (Rs) from steady-state voltage responses. Rheobase was determined using a slow current ramp (-200 to 700 pA at 0.9 pA/ms). No DC current was applied to adjust the resting membrane potential (RMP). Bridge balance and pipette capacitance compensation were adjusted throughout recordings. Data were discarded if R_s_ exceeded 20 MΩ or changed by >20% during the experiment.

#### 2.2.3 EPSP summation measurements

Spontaneous EPSPs (sEPSPs) were recorded in current-clamp mode at 0 pA holding current. For evoked EPSPs, a tungsten bipolar stimulating electrode (FHC Inc.) was positioned in layer 1 (200 μm from the soma of the recorded cell). Electrical stimuli (0.1-0.3 mA; 0.2 ms duration) were delivered via a WPI A395 stimulus isolator at 15-30% of the maximum stimulation intensity, evoking EPSPs of 10 ± 5 mV amplitude. Trains of five stimuli at 50 Hz were used to assess EPSP summation.

Signals were low-pass filtered at 10 kHz and digitized at 50 kHz using a Multiclamp 700A amplifier, a National Instruments PCIe-6323 board and controlled with WinWCP v5.8 software (J. Dempster, University of Strathclyde; https://github.com/johndempster/WinWCPXE).

#### 2.2.4 Voltage Clamp Recordings

Voltage-clamp recordings were obtained from L2/3 pyramidal neurons using borosilicate pipettes (4-6 MΩ) filled with an internal solution containing (in mM): 120 K-gluconate, 10 KCl, 8 NaCl, 10 HEPES, 0.5 EGTA, 4 Mg-ATP, 0.3 Na-GTP, 10 phosphocreatine, and 1 QX-314 (pH 7.2; 295-305 mOsm/kg). R_s_ was monitored and compensated by 80%. Pipette and whole-cell capacitances were fully compensated. Data were discarded if R_s_ exceeded 20 MΩ or changed by >20% during the experiment.

Cells were held at -80 mV in ACSF containing pharmacological blockers to isolate specific currents. The external solution contained: 0.5 μM tetrodotoxin (TTX), 5 mM tetraethylammonium (TEA), 5 mM 4-aminopyridine (4-AP),10 μM CNQX, 100 μM DL-AP5, and 50 μM picrotoxin to isolate I_h_. To isolate Ba^2+^-sensitive inwardly rectifying potassium current, we use the same solution with 10 μM ZD7288 added.

Voltage protocols consisted of step commands from -130 to-40 mV in 10 mV increments (800 ms duration), followed by a voltage ramp from -130 to -40 mV over 500 ms at 1 Hz. The liquid junction potential (LJP) was calculated as 16.2 mV using the stationary Nernst-Planck equation and LJPcalc (https://swharden.com/LJPcalc) and were not subtracted. Reversal potentials were calculated as the difference between control current and Ba^2+^-sensitive current using the voltage ramp protocols.

#### 2.3 I_h_ and inward rectifying potassium current inhibition

To assess the contributions of HCN and Ba^2+^-sensitive inwardly rectifying potassium current, 10 μM ZD7288 and 60 μM BaCl_2_ were individually applied to the bath solution in separate experimental sets to selectively inhibit Ih and Ba^2+^-sensitive inwardly rectifying potassium current, respectively. Inhibitors were perfused at the same concentrations during both current clamp and voltage clamp recordings.

### 2.4 Data analysis

Electrophysiological data were analyzed using Clampfit v10.3 (Molecular Devices) and Igor Pro v6.37 (WaveMetrics) with the NeuroMatic plugin (Rothman & Silver, 2018). Spontaneous EPSPs were detected and quantified using the miniML Python module (O’Neill et al., 2025), using identical detection parameters across genotypes and drug conditions. Rheobase was defined as the minimum depolarizing current required to evoke a single action potential and was quantified from ramp protocols (−200 to 700 pA at 0.9 pA/ms) to improve precision relative to step-based estimates.

Data are presented as the mean ± 95% confidence interval (95% C.I.), as indicated. In figures, individual data points are plotted alongside group summaries; where shown, the “mean difference” panel represents the paired or unpaired difference between conditions or genotypes (Gardner–Altman estimation plots).

Normality was assessed using the Shapiro-Wilk test. For paired comparisons, normally distributed data were analyzed using paired Student’s t-tests and non-parametric data using Wilcoxon signed-rank tests. For unpaired comparisons, normally distributed data were analyzed using unpaired Student’s t-tests (Welch’s correction applied when variances were unequal) and non-parametric data using Mann–Whitney U tests. For analyses involving multiple current steps and/or drug conditions, two-way ANOVA was used as indicated in the Results, with appropriate post hoc multiple-comparison testing when applicable. Statistical significance was set at p < 0.05. All statistical analyses and data visualization were performed using GraphPad Prism v8 (GraphPad Software, USA). Sample sizes are reported as n (neurons) and N (mice).

## 3. RESULTS

### 3.1 Ba^2+^-sensitive potassium conductance underlies hypoexcitability of L2/3 primary auditory cortex pyramidal neurons in Fmr1-KO mice

First, we investigated passive membrane properties of layer 2/3 (L2/3) primary auditory cortex pyramidal neurons in acute slices from wild-type (WT) and Fmr1-KO mice. Compared with WT, pyramidal neurons from Fmr1-KO mice exhibited a more hyperpolarized resting membrane potential (RMP; WT: -78.3 ± 6.4 mV; Fmr1-KO: -82.0 ± 6.8 mV; p = 0.0457; Fig. 1A,B) and lower input resistance (R_in_; WT: 172.7 ± 34.7 MΩ; Fmr1-KO: 124.5 ± 33.2 MΩ, p < 0.001; Fig. 1C), consistent with an elevated basal conductance. Bath application of BaCl_2_ (60 μM) depolarized the RMP and increased R_in_ in Fmr1-KO neurons (ΔRMP: 4.8 mV; ΔR_in_: 66.6 MΩ; p < 0.001; Fig. 1A-C), shifting both measures toward WT values; effects in WT were smaller (ΔRMP: 1.5 mV; ΔR_in_: 51.1 MΩ; Fig. 1A-C).

**Figure 1.**
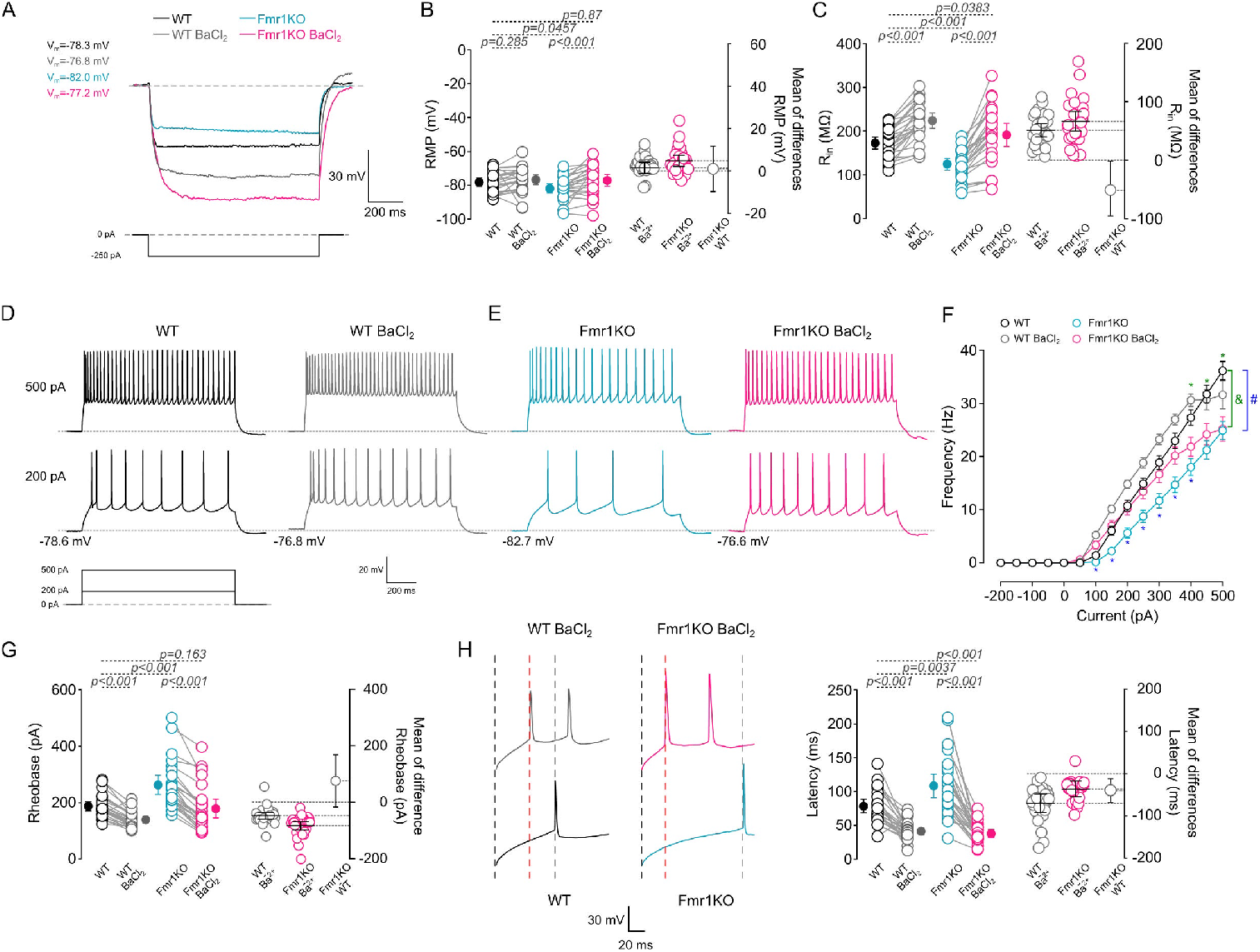
Ba^2+^-sensitive inwardly rectifying potassium conductance constrains passive properties and firing output in L2/3 primary auditory cortex pyramidal neurons. (A) Representative voltage responses to a 250 pA current step recorded from WT (black) and Fmr1-KO (light blue) L2/3 pyramidal neurons before and after bath application of BaCl_2_ (60 μM; WT: gray; Fmr1-KO: pink). (BC) Summary quantification of resting membrane potential (RMP) and input resistance (R_in_) before and after BaCl_2_. (DE) Representative firing patterns at rheobase and during 200 and 500 pA depolarizing steps, before and after BaCl_2_. (F) Inputoutput (I/O) relationships for WT and Fmr1-KO neurons under control conditions and following BaCl_2_. (G) Rheobase measured from currentramp protocols. (H) Representative first action potential at rheobase and summary quantification of firstspike latency. Data are mean ± 95% C.I.; individual data points are shown. WT: n = 26 neurons, N = 8 mice; Fmr1-KO: n = 24 neurons, N = 8 mice.

Next, we tested whether inhibiting Ba^2+^-sensitive inwardly rectifying potassium current alters firing output. At baseline, Fmr1-KO neurons fired fewer spikes across 200-500 pA steps (two-way ANOVA, p < 0.001; Fig. 1D-F) and exhibited an increased rheobase (WT: 187.2 ± 39.8 pA; Fmr1-KO: 262.9 ± 84.6 pA; p < 0.001; Fig. 1G) together with a longer latency to the first action potential (WT: 78.4 ± 24.4 ms; Fmr1-KO: 108.4 ± 43.3 ms; p = 0.0037; Fig. 1H). BaCl_2_ reduced rheobase in Fmr1-KO neurons (Δ = -84.1 pA; Fig. 1G) and restored the input-output relationship toward WT-like ranges (after BaCl_2_, WT vs Fmr1-KO: one-way ANOVA, p = 0.82; Fig. 1F); first-spike latency also decreased in Fmr1-KO neurons (Fig. 1H).

Because changes in RMP and R_in_ can influence recruitment of voltage-dependent conductances involved in spike initiation, we analyzed the waveform of the first action potential at rheobase. Fmr1-KO neurons exhibited a slightly more hyperpolarized spike threshold (WT: -36.1 mV; Fmr1-KO:-38.4 mV; p = 0.0099; Fig. 2A,B) and increased amplitude (WT: 103.5 mV; Fmr1-KO: 110.4 mV; p = 0.022; Fig. 2A,C),with no significant change in spike duration (FWHM; WT: 1.41 ms; Fmr1-KO: 1.32 ms; p = 0.151; Fig. 2A,D). In Fmr1-KO neurons, BaCl_2_ depolarized the spike threshold (Δ = 3.1 mV; p < 0.001; Fig. 2B), reduced spike amplitude (Δ = -7.4 mV; p < 0.001; Fig. 2C), and increased spike duration (ΔFWHM = 1.1 ms; p < 0.001; Fig. 2D), indicating that Ba^2+^-sensitive potassium conductance shapes both passive properties and spike waveform. Together, these results support the conclusion that enhanced Ba^2+^-sensitive potassium conductance promotes a hyperpolarized, low-R_in_, hypoexcitable state in Fmr1-KO L2/3 primary auditory cortex neurons that is reversible with Ba^2+^.

**Figure 2.**
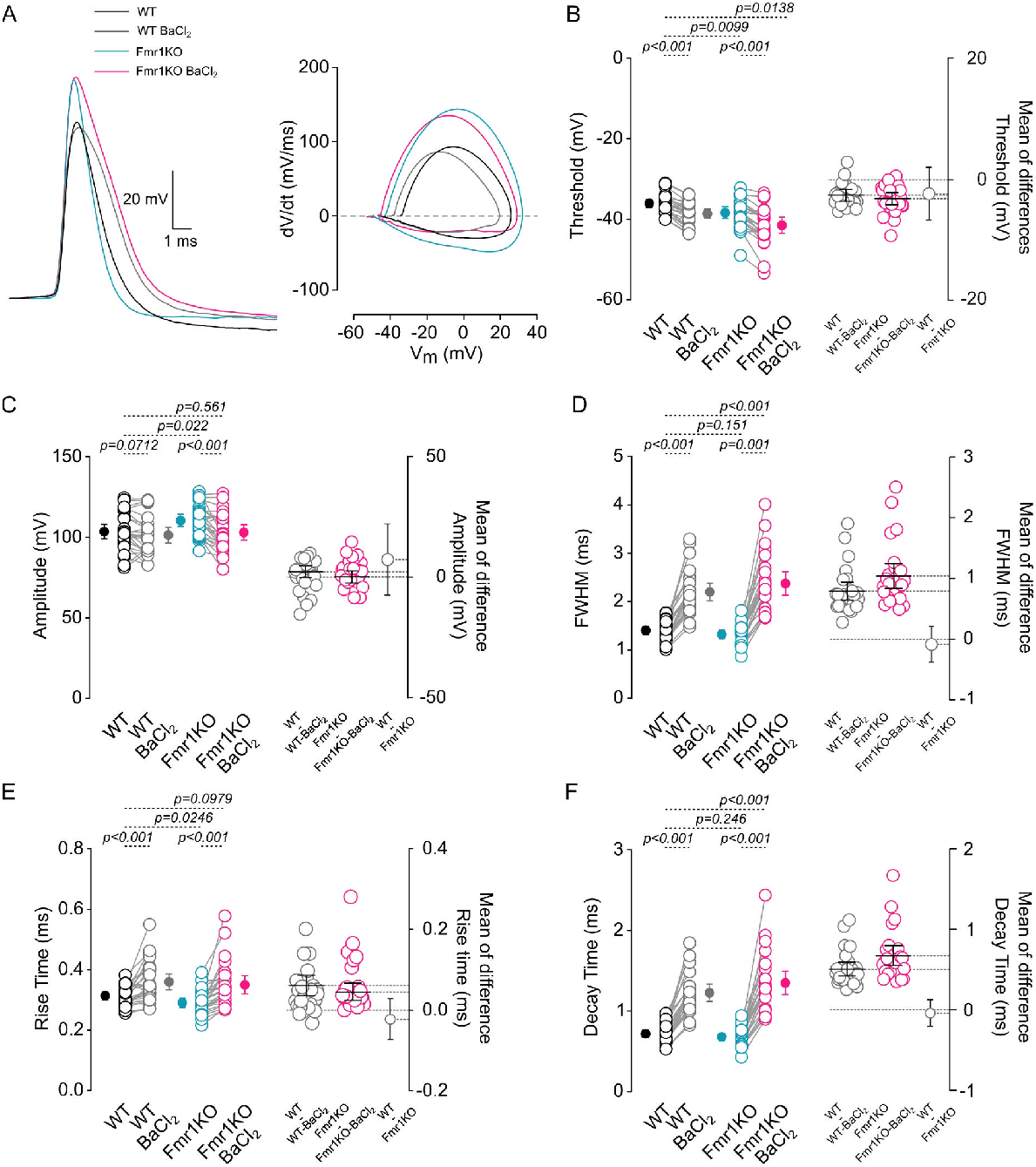
Ba^2+^ application modifies action potential waveform in Fmr1-KO L2/3 pri mary auditory cortex pyramidal neurons. (A) Representative first action potentials (APs) elicited at rheobase and corresponding phase plots (dV/dt vs. V) from WT and Fmr1-KO neurons, shown before and after BaCl_2_ (60 μM). For visualization, all APs are displayed from the same baseline and aligned to AP onset. (B–F) Summary quantification of AP threshold, AP amplitude, AP duration (full width at half maximum; FWHM), rise time, and decay time under control conditions and following BaCl_2_. Data are mean ± 95% C.I.; individual data points are shown. WT: n = 26 neurons, N = 8 mice; Fmr1-KO: n = 24 neurons, N = 8 mice.

### 3.2 Synaptic input and integration are elevated, but independent of Ba^2+^-sensitive inward rectifying potassium conductance

Kir channels influence synaptic integration by modulating the response to synaptic inputs and influencing transmitter release, so we next asked whether Ba^2+^-sensitive potassium conductance alters synaptic activity in both genotypes. We first quantified spontaneous excitatory postsynaptic potential (sEPSP) frequency at rest and found that Fmr1-KO neurons exhibited higher sEPSP frequency than WT (WT: 3.56 ± 0.89 Hz; Fmr1-KO: 4.41 ± 1.06 Hz; p = 0.003; Fig. 3A,B). BaCl_2_ (60 μM) did not significantly change sEPSP frequency in either genotype (WT Δ: 0.09 Hz; Fmr1-KO Δ: 0.04 Hz; all p > 0.05; Fig. 3A,B), arguing against a strong presynaptic effect under these conditions.

**Figure 3.**
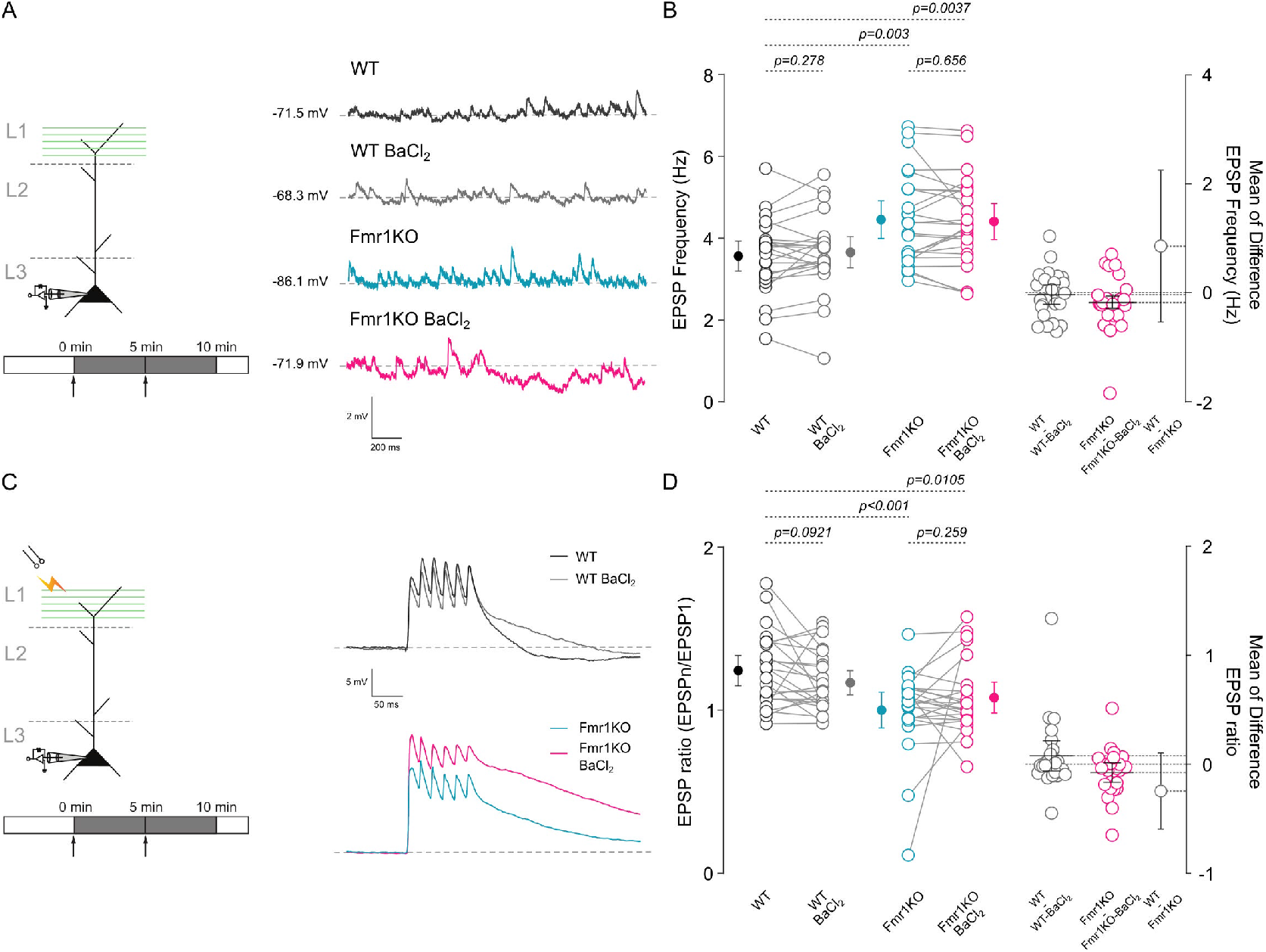
Elevated synaptic activity and reduced EPSP summation in Fmr1-KO neurons are not significantly altered by Ba^2+^. (A) Representative spontaneous excitatory postsynaptic potentials (sEPSPs) recorded in current clamp at 0 pA holding current from WT and Fmr1-KO neurons before and after BaCl_2_ (60 μM). (B) Summary of sEPSP frequency under control conditions and after BaCl_2_. (C) Representative EPSP summation responses evoked by 50 Hz extracellular stimulation in layer 1 (L1) of the pri mary auditory cortex. (D) Summary quantification of EPSP summation ratio (as defined in Methods) before and after BaCl_2_. Data are mean ± 95% C.I.; individual data points are shown. WT: n = 25 neurons, N = 8 mice; Fmr1-KO: n = 24 neurons, N = 8 mice.

We then assessed synaptic integration using EPSP summation during 50 Hz layer-1 stimulation (Methods). Fmr1-KO neurons exhibited a lower EPSP summation ratio than WT (WT: 1.2 ± 0.2; Fmr1-KO: 1.0 ± 0.3; p < 0.001; Fig. 3C,D). BaCl_2_ did not significantly alter summation in either genotype (Δ ratio: WT = 0.07; Fmr1-KO = 0.08; all p > 0.05; Fig. 3C,D). Thus, while basal excitatory synaptic activity is elevated in Fmr1-KO L2/3 neurons, Ba^2+^-sensitive potassium conductance primarily constrains intrinsic excitability rather than measurably changing sEPSP frequency or short-train EPSP summation.

### 3.3 Increased functional Ba^2+^-sensitive inward rectifying potassium current in L2/3 pyramidal neurons of the primary auditory cortex from Fmr1-KO mice

Because Ba^2+^-sensitive conductance strongly influenced excitability in Fmr1-KO neurons, we next quantified the underlying subthreshold current in voltage clamp (Methods). In both genotypes we observed a Ba^2+^-sensitive inwardly rectifying current (Fig. 4A,B). However, the Ba^2+^-sensitive current amplitude at -130 mV was larger in Fmr1-KO neurons (WT: -218.2 ± 70.2 pA; Fmr1-KO: -307.3 ± 89.2 pA; p = 0.0302; Fig. 4B,C), and the percent Ba^2+^ block at -130 mV was also greater (WT: 59.6 ± 12.0%; Fmr1-KO: 76.9 ± 7.9%; p = 0.0045; Fig. 4C). Ba^2+^-subtracted I-V relationships showed comparable rectification, and reversal potentials were not significantly different between genotypes (WT: -89.6 ± 2.4 mV; Fmr1-KO: -82.9 ± 16.3 mV; p = 0.126). Together, these data indicate an increased functional contribution of Ba^2+^-sensitive potassium conductance in Fmr1-KO L2/3 primary auditory cortex pyramidal neurons, consistent with their hyperpolarized RMP, reduced R_in_, and reduced firing output.

**Figure 4.**
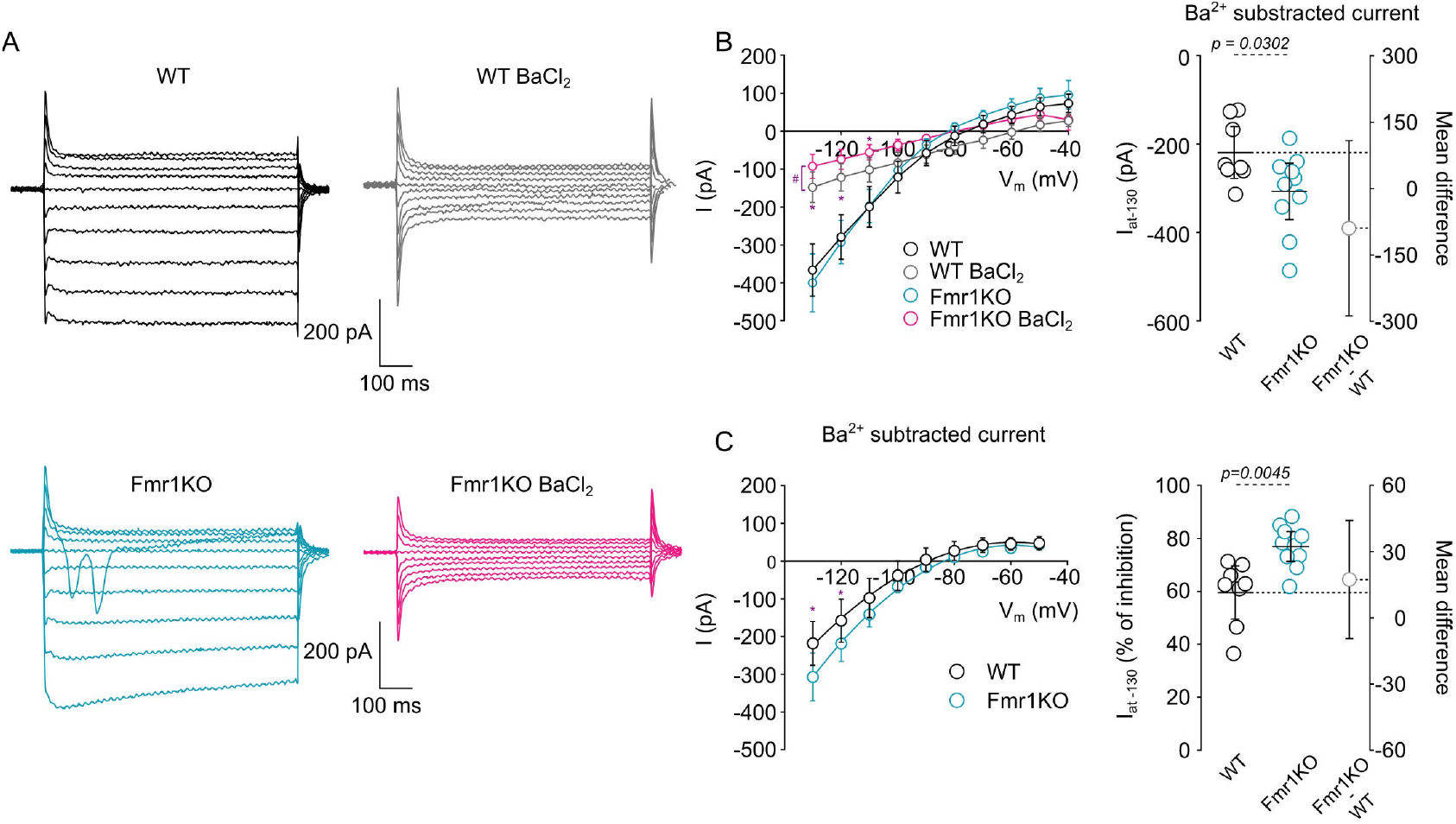
Increased Ba^2+^-sensitive inwardly rectifying potassium current in Fmr1-KO L2/3 primary auditory cortex pyramidal neurons. (A) Representative Ba^2+^-sensitive inwardly rectifying potassium current recorded in voltage clamp from WT and Fmr1-KO neurons before and after BaCl_2_ (60 μM), and the corresponding Ba^2+^-sensitive (subtracted) component. (B) Currentvoltage (IV) relationships before and after BaCl_2_. (CD) Summary quantification of Ba^2+^-sensitive current amplitude at −130 mV and the fraction of current blocked by Ba^2+^ at 130 mV. Data are mean ± 95% C.I.; individual data points are shown. n = 8 neurons, N = 4 mice per genotype.

### 3.4 I_h_ makes a secondary, inputfiltering contribution with genotypespecific effects

We next examined I_h_, mediated by HCN channels, as a second major contributor to subthreshold conductance. Blocking I_h_ with ZD7288 (10 μM) hyperpolarized RMP in both genotypes (ΔWT: -4.7 mV; ΔFmr1-KO: -3.3 mV; all p < 0.05; Fig. 5A,B) and increased R_in_ (ΔWT: 8.0 MΩ, p = 0.0349; ΔFmr1-KO: 12.1 MΩ, p < 0.001; Fig. 5C) while abolishing sag (pre-ZD7288: WT 1.8 mV, Fmr1-KO 1.5 mV, p = 0.407; post-ZD7288: WT 1.0 mV, p = 0.0257; Fmr1-KO 1.1 mV, p = 0.0061; Fig. 5D). Notably, ZD7288 did not shift passive properties in Fmr1-KO neurons toward WT values to the same extent observed with BaCl_2_, suggesting that I_h_ is not the primary determinant of the Fmr1-KO hypoexcitable state.

**Figure 5.**
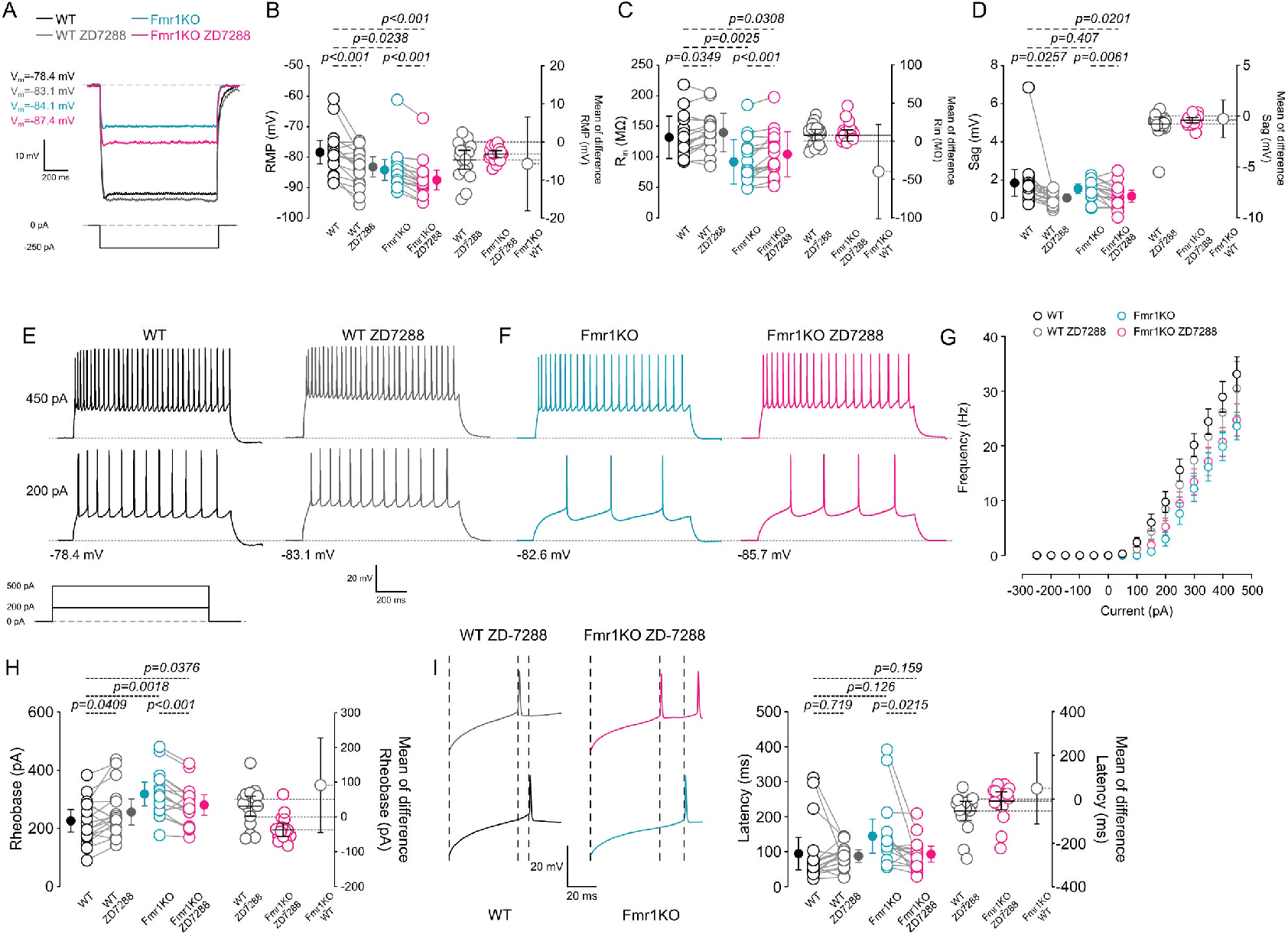
I_h_ inhibition alters passive properties and produces genotypedependent effects on intrinsic excitability. (A) Representative voltage responses to a 250 pA current step recorded from WT (black) and Fmr1-KO (light blue) neuronsbefore and after ZD7288 (10 μM; WT: gray; Fmr1-KO: pink). (BD) Summary quantification of resting membrane potential (RMP), input resistance (R_in_), and sag amplitude before and after ZD7288. (EF) Representative firing patterns at rheobase and during 200 and 500 pA depolarizing steps, before and after ZD7288. (G) Inputoutput (I/O) relationships for WT and Fmr1-KO neurons under control conditions and following ZD7288. (H) Rheobase measured from currentramp protocols. (I) Representative first AP at rheobase and summary quantification of firstspike latency. Data are mean ± 95% C.I.; individual data points are shown. WT: n = 20 neurons, N = 7 mice; Fmr1-KO: n = 17 neurons, N = 7 mice.

To test the impact of I_h_ on intrinsic excitability, we repeated current-step protocols in both genotypes (Methods). ZD7288 increased rheobase in WT neurons (Δ = 29.8 pA; p = 0.0409; Fig. 5E-H) but decreased rheobase in Fmr1-KO neurons (Δ = -37.5 pA; p < 0.001; Fig. 5E-H), yielding a significant genotype-drug interaction for firing output (two-way ANOVA interaction, p < 0.001; Fig. 5E-G). First-spike latency decreased in both genotypes (Fig. 5I). ZD7288 increased AP duration (FWHM) in both WT and Fmr1-KO neurons (ΔWT: 0.15 ms, p < 0.001; ΔFmr1-KO: 0.11 ms, p = 0.0042; Fig. 6A,D), whereas AP threshold and amplitude were not significantly changed (Fig. 6A-C).

**Figure 6.**
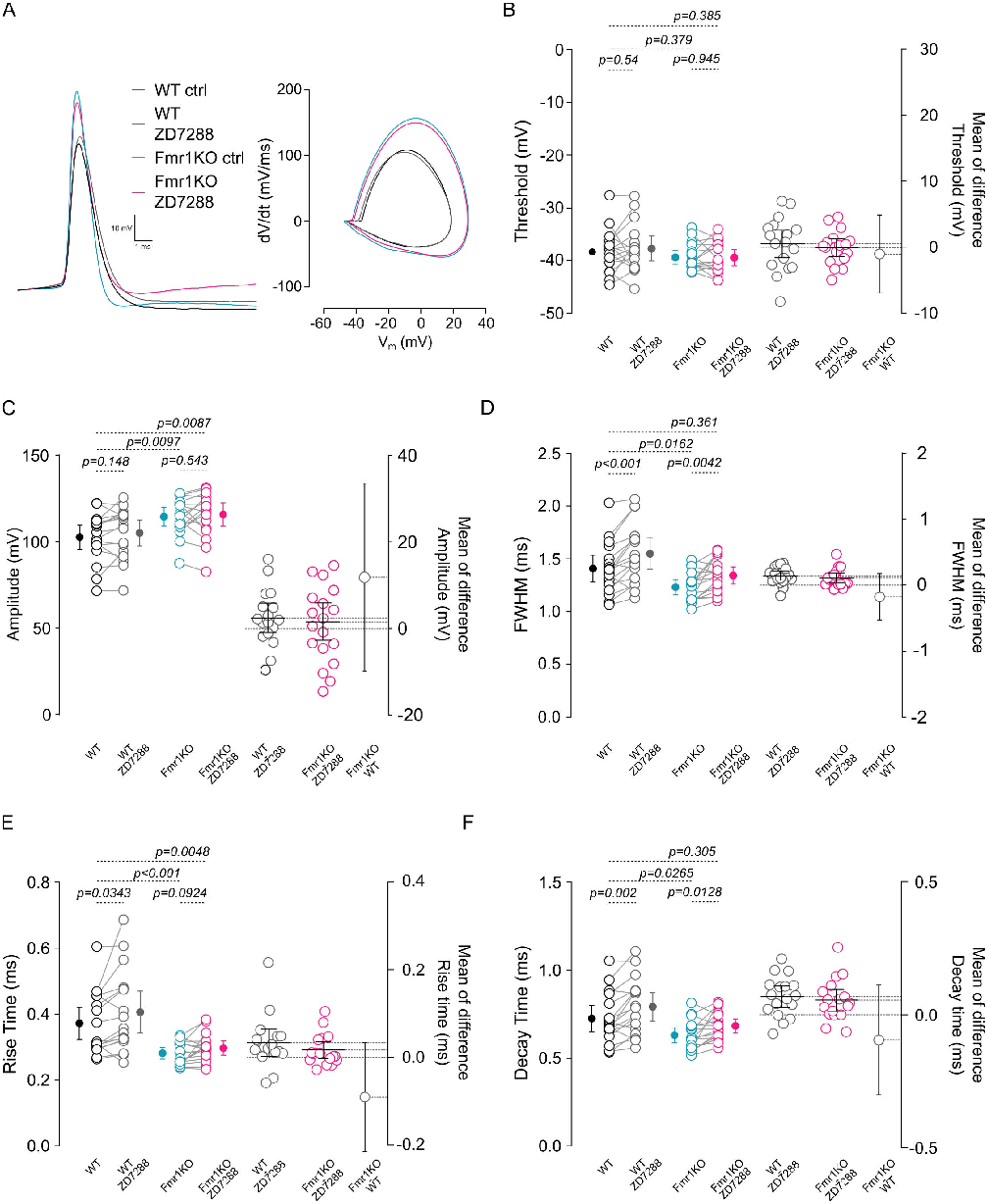
I_h_ inhibition increases action potential duration in WT and Fmr1-KO L2/3 primary auditory cortex pyramidal neurons. Representative first action potentials (APs) elicited at rheobase and corresponding phase plots (dV/dt vs. V) from WT and Fmr1-KO neurons, shown before and after ZD7288 (10 μM). For visualization, all APs are displayed from the same baseline and aligned to AP onset. (BF) Summary quantification of AP threshold, AP amplitude, AP duration (FWHM), rise time, and decay time under control conditions and following ZD7288. Data are mean ± 95% C.I.; individual data points are shown. WT: n = 20 neurons, N = 7 mice; Fmr1 KO: n = 17 neurons, N = 7 mice.

Because I_h_ is prominent in dendritic compartments and shapes synaptic integration, we next tested how I_h_ inhibition affects synaptic transmission. ZD7288 increased sEPSP frequency in both genotypes (ΔWT: 0.28 Hz, p = 0.029; ΔFmr1-KO: 1.01 Hz, p = 0.0015; Fig. 7A,B) and enhanced EPSP summation selectively in Fmr1-KO neurons (Δ ratio: 0.28, p = 0.0086; Fig. 7C,D). Together, these findings suggest that I_h_ contributes to dendritic filtering and produces genotype-dependent shifts in excitability and synaptic integration, whereas Ba^2+^-sensitive inward rectifying potassium current remains the dominant factor setting the Fmr1-KO hypoexcitable state.

**Figure 7.**
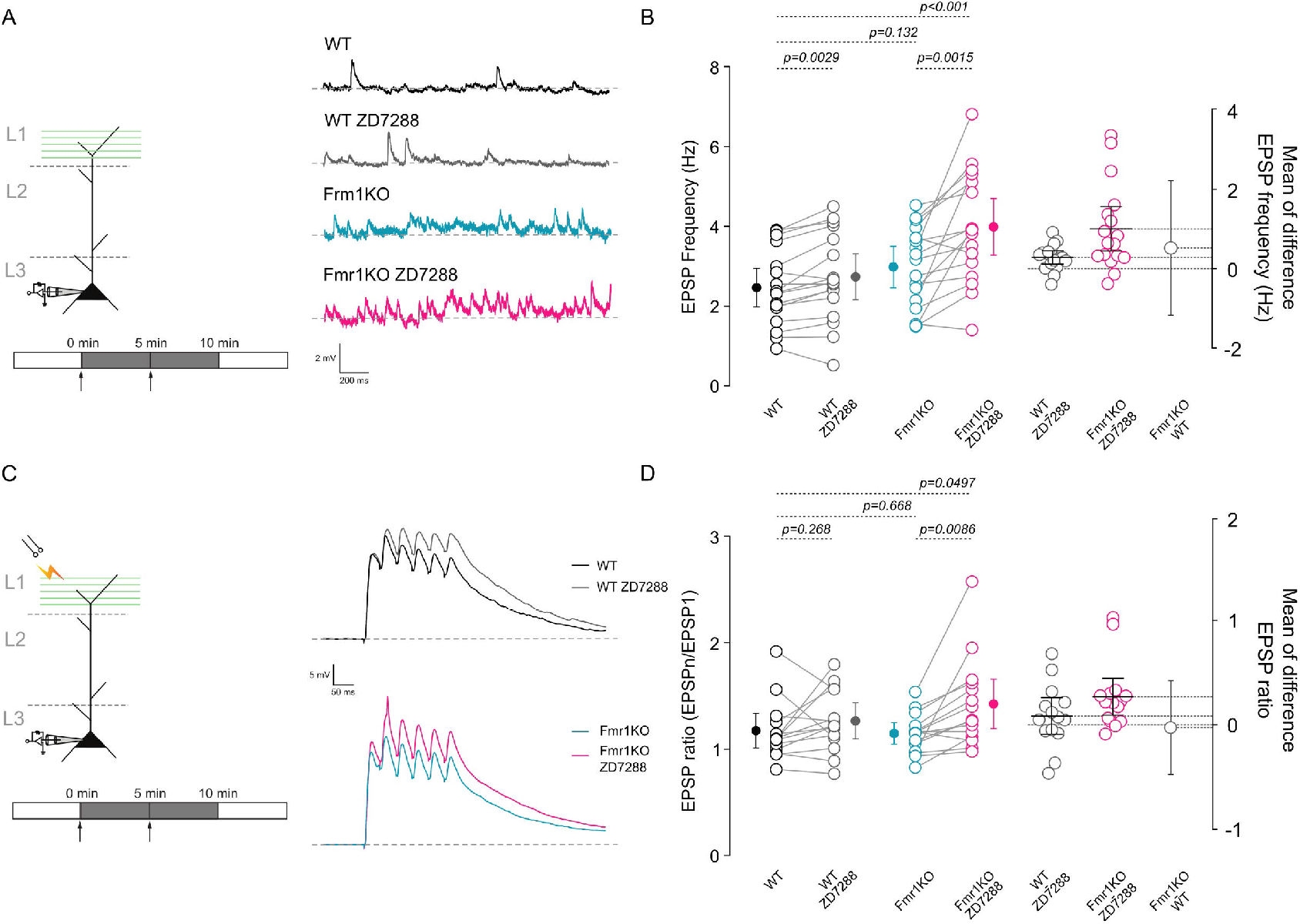
I_h_ inhibition increases synaptic activity and enhances EPSP summation preferentially in Fmr1-KO neurons. (A) Representative sEPSPs recorded in current clamp at 0 pA holding current from WT and Fmr1-KO neurons before and after ZD7288 (10 μM). (B) Summary of sEPSP frequency under control conditions and after ZD7288. (C) Representative EPSP summation responses evoked by 50 Hz extracellular stimulation in layer 1 of the primary auditory cortex. (D) Summary quantification of EPSP summation ratio before and after ZD7288. Data are mean ± 95% C.I.; individual data points are shown. WT: n = 17 neurons, N = 7 mice; Fmr1-KO: n = 17 neurons, N = 7 mice.

## 4. DISCUSSION

In this study, we demonstrate reduced intrinsic excitability of layer 2/3 (L2/3) pyramidal neurons in the primary auditory cortex of Fmr1-KO mice. Despite elevated excitatory synaptic activity onto these neurons, their firing output was reduced, consistent with a compensatory shift toward a hypoexcitable state. This phenotype was accompanied by a hyperpolarized resting membrane potential (RMP), reduced input resistance, increased rheobase, and prolonged first-spike latency. Pharmacological inhibition with BaCl_2_ restored key excitability measures toward WT levels, supporting the interpretation that an enhanced Ba^2+^-sensitive potassium conductance constrains intrinsic excitability in Fmr1-KO L2/3 primary auditory cortex pyramidal neurons. In contrast, I_h_ inhibition produced more modest changes in passive properties and yielded genotype-dependent effects on firing and synaptic integration, consistent with a secondary role in input filtering rather than setting basal conductance.

### 4.1 Reduced excitability of L2/3 pyramidal neurons of the primary auditory cortex in Fmr1-KO mice

Our experiments revealed elevated excitatory synaptic activity accompanied by reduced intrinsic excitability in L2/3 primary auditory cortex pyramidal neurons from Fmr1-KO mice. These findings are consistent with homeostatic compensation, in which neurons reduce gain to stabilize output in the face of increased excitatory drive. Across brain regions, Fmr1-KO circuits exhibit altered intrinsic excitability, often linked to changes in ion channel expression, modulation, or subcellular distribution that shape passive properties, action potential initiation and waveform, and synaptic integration (Contractor et al., 2015; Booker et al., 2019; McCullagh et al., 2020; Zhan et al., 2020; Susco et al., 2020; Deng & Klyachko, 2021; Ordemann et al., 2021; Bülow et al., 2022; Brandalise et al., 2023). Collectively, these studies support the idea that compensatory mechanisms may emerge to stabilize network activity.

FMRP modulates ion channel function through multiple mechanisms, including regulation of expression, mRNA transport, and direct protein-protein interactions (El-Hassar et al., 2019; Deng & Klyachko, 2021; Ordemann et al., 2021; Bülow et al., 2022). For example, in hippocampal pyramidal neurons, loss of FMRP can reduce expression of Kv3.1 and Kv4.2 (Gross et al., 2011). In cortical neuron cultures, disruption of the FMRP-β4 subunit interaction at BK channels increases BK open probability (Deng & Klyachko, 2021; Mitchell et al., 2023), which can shorten action potentials and alter EPSP integration, thereby modifying excitability (Deng & Klyachko, 2016; Mitchell et al., 2023; Ferraguto et al., 2023).

In our recordings, Fmr1-KO pyramidal neurons exhibited a hyperpolarized RMP, reduced input resistance, and increased rheobase, consistent with a change in conductance state that limits firing in response to weak inputs. The conductance states can suppress spontaneous firing in hyperexcitable environments and help stabilize firing rates (Carrasquillo et al., 2012; Kourrich et al., 2015; Combe et al., 2018; Debanne et al., 2019).

The basal conductance states arise from increased subthreshold conductances. I_h_ and Kir channels both contribute to baseline conductance but exert opposing effects: I_h_ depolarizes the membrane and can promote subthreshold resonance, whereas Kir channels hyperpolarize and stabilize the RMP (Day et al., 2005a; González et al., 2012; Bataveljic et al., 2024). In our experiments, the magnitude of Ba^2+^-sensitive effects on passive properties and firing, together with the absence of a strong genotype-dependent shift in I_h_-dependent baseline conductance, supports the conclusion that Ba^2+^-sensitive potassium currents are the principal contributors to the Fmr1-KO hypoexcitable state.

It is worth noting that the reduced basal intrinsic excitability of L2/3 pyramidal neurons in Fmr1-KO mice suggests that these neurons reside in a hypoexcitable state, which contrasts with what has been reported in other cortical regions. For example, hyperexcitability of pyramidal neurons across multiple layers has been described in the visual, entorhinal, piriform, somatosensory, and medial prefrontal cortices, largely driven by changes in active membrane properties, such as a lower action potential (AP) threshold and altered expression of voltage-gated Na^+^ and K^+^ channels, that modify AP onset, threshold, and waveform, thereby increasing firing frequency with little or no change in passive properties (Zhang et al., 2016; Deng & Klyachko, 2016b; Liu et al., 2022; Arancibia et al., 2025).

Although we observed qualitatively similar changes in active properties in the recorded L2/3 pyramidal neurons from Fmr1-KO mice compared with WT, these effects were not strong enough to overcome the hypoexcitable state imposed by changes in passive properties. This hypoexcitable, high-conductance state may also be shaped by impaired excitation/ inhibition (E/I) balance. In the aforementioned cortical areas, reduced GABAergic tone has been associated with decreases in parvalbumin (PV) interneuron number and intrinsic excitability (Wen et al., 2018). In our recordings from AC1, excitatory drive was also increased, consistent with findings in those regions. However, PV interneuron density in AC1 has been reported to be reduced during development in Fmr1-KO mice and to recover to WT levels by approximately P30–P40 (Wen et al., 2018). Because the mice used here fall within this recovery window, it is unlikely that the observed hypoexcitable state is explained by an increased basal inhibitory tone.

### 4.2 Augmented Ba^2+^-sensitive potassium current in L2/3 pyramidal neurons of the primary auditory cortex in Fmr1-KO mice

Our results indicate that L2/3 primary auditory cortex pyramidal neurons in Fmr1-KO mice exhibit enhanced Ba^2+^-sensitive potassium conductance that reduces input resistance and hyperpolarizes the RMP, thereby lowering neuronal gain and increasing rheobase. This places Fmr1-KO neurons in a stabilized, low-excitability regime that requires stronger depolarizing drive to reach the spike threshold.

Kir channels can establish high-conductance states and thereby constrain excitability (Day et al., 2005a; Johnson-Venkatesh et al., 2015). Two major Kir families are expressed in the CNS: Kir2.x channels, which are strongly inwardly rectifying and activated by hyperpolarization, and Kir3.x (GIRK) channels, which are gated by G-protein-coupled receptors (Hibino et al., 2010; Lüscher & Slesinger, 2010). Although direct regulation of neuronal Kir2.x or Kir3.x channels by FMRP has not been established, FMRP broadly regulates ion channel expression and function via post-transcriptional and protein-protein mechanisms (Contractor et al., 2015; Ferron, 2016; McCullagh et al., 2020; Deng & Klyachko, 2021), raising the possibility that loss of FMRP increases Kir channel expression or functional availability. Future experiments using single-cell RT-PCR, immunofluorescence, or targeted pharmacology will be required to determine which Kir subtypes are altered and whether changes reflect expression, localization, or modulation.

Consistent with this interpretation, voltage-clamp recordings revealed a larger Ba^2+^-sensitive inwardly rectifying current between -130 to -110 mV in Fmr1-KO neurons and a greater fraction of Ba^2+^-sensitive current, while Ba^2+^-subtracted I-V relationships showed similar rectification and no significant genotype difference in reversal potential. Together, these findings support an increased functional contribution of Ba^2+^-sensitive potassium conductance in Fmr1-KO neurons. Nevertheless, we cannot fully exclude off-target effects of Ba^2+^. For example, even relatively low concentrations of Ba^2+^ have been reported to inhibit A-type potassium currents under some conditions (Kehl et al., 2013), which could influence excitability.

Multiple studies report altered A-type potassium currents in Fmr1-KO models (Gross et al., 2011; Lee et al., 2011; Routh et al., 2013; Zhan et al., 2020; Deng & Klyachko, 2021; Ordemann et al., 2021), and A-type channels (notably Kv4.2) can delay spike onset and increase rheobase by opposing depolarization near threshold (Sacco & Tempia, 2002; Carrasquillo et al., 2012; Xie et al., 2014; Xiao et al., 2014; Rathour et al., 2016). In our dataset, the strongest genotype differences aligned with passive properties (RMP and Rin) and firing output, while BaCl_2_ also modified spike waveform. Thus, it is plausible that Ba^2+^-sensitive inwardly rectifying potassium conductance is a primary driver of the conductance state, with additional contributions from other K^+^ conductances (including A-type currents) that warrant direct testing in future work.

In summary, L2/3 pyramidal neurons in the primary auditory cortex of Fmr1-KO mice receive elevated excitatory synaptic input yet exhibit reduced intrinsic excitability consistent with an increase in basal conductance. This compensation depends strongly on Ba^2+^-sensitive inward rectifying potassium currents with Kir-like properties, whose inhibition depolarizes the RMP, increases input resistance, and restores firing output without measurably changing sEPSP frequency or short-train EPSP summation. Such intrinsic regulation may stabilize cortical output but alter sensory integration, contributing to abnormal auditory processing in Fragile X syndrome.

### 4.3 Proposed model of Ba^2+^-sensitive inwardly rectifying potassium currentdependent homeostatic hypoexcitability

Together, our findings support a model in which L2/3 primary auditory cortex pyramidal neurons in Fmr1-KO mice engage a Ba^2+^-sensitive inwardly rectifying potassium conductance that acts as a homeostatic brake against excessive excitatory drive. Enhanced Ba^2+^-sensitive potassium conductance hyperpolarizes the membrane, reduces input resistance, and decreases neuronal gain, thereby stabilizing firing output despite increased upstream excitation. While this intrinsic adaptation may preserve network stability, it is also likely to alter input integration and timing, providing a plausible cellular substrate for disrupted auditory processing associated with Fragile X syndrome.

## Supplementary information

Tables with all the electrophysiological parameters are included in the supplementary tables file.

## Ethics approval

All animal experiments were approved by the Ethics Committee of the University of Santiago de Chile (protocol 034.2023).

## Consent for publication

All authors consent to publication.

## Availability of data and materials

All data generated or analyzed during this study are included in this published article.

## Author contributions

C.M. and E.L.-S. conceived the study. C.M., D.R., and E.L.-S. performed experiments and analyzed data. C.M., D.R., C.C.-d.R., and E.L.-S. interpreted the results. C.C.-d.R. provided reagents. C.M., D.R., and E.L.-S. drafted the manuscript, and all authors revised and approved the final version.

