## Supplementary tables for "Enhanced Ba^2+^-sensitive inward rectifying potassium conductance reduces intrinsic excitability of layer 2/3 pyramidal neurons in the primary auditory cortex of Fmr1 knockout mice"

Table 1. Intrinsic excitability parameteros ($BaCl_2$)

| | WT | SD | WT $BaCl_2$ | SD | Statistical test | P-value | |
|---|---|---|---|---|---|---|---|
| n | | | | 26 | | | |
| RMP (mV) | -78.3 | 6.3 | -76.8 | 7.2 | Paired t test | 0.285 | |
| Rin (MΩ) | 172.7 | 34.7 | 223.8 | 43.4 | Paired t test | <0.001 | * |
| Tau (ms) | 14.8 | 4.8 | 36.0 | 13.7 | Wilcoxon test | <0.001 | * |
| Rheobase (pA) | 187.2 | 39.8 | 139.3 | 27.8 | Paired t test | <0.001 | * |
| AP latency (ms) | 78.4 | 24.4 | 41.4 | 13.3 | Paired t test | <0.001 | * |
| AP threshold (mV) | -36.1 | 2.4 | -38.7 | 2.9 | Paired t test | <0.001 | * |
| AP amplitude (mV) | 103.5 | 11.7 | 101.1 | 11.8 | Paired t test | 0.0712 | |
| FWHM (ms) | 1.41 | 0.21 | 2.20 | 0.45 | Wilcoxon test | <0.001 | * |
| Rise time (ms) | 0.31 | 0.03 | 0.36 | 0.07 | Paired t test | <0.001 | * |
| Decay time (ms) | 0.73 | 0.12 | 1.23 | 0.27 | Wilcoxon test | <0.001 | * |

| | Fmr1KO | SD | Fmr1KO $BaCl_2$ | SD | Statistical test | P-value | |
|---|---|---|---|---|---|---|---|
| n | | | | 26 | | | |
| RMP (mV) | -82.0 | 6.8 | -77.2 | 8.9 | Paired t test | <0.001 | * |
| Rin (MΩ) | 124.5 | 33.2 | 191.1 | 65.0 | Paired t test | <0.001 | * |
| Tau (ms) | 12.0 | 4.2 | 34.1 | 14.4 | Wilcoxon test | <0.001 | * |
| Rheobase (pA) | 262.9 | 84.6 | 178.8 | 81.1 | Wilcoxon test | <0.001 | * |
| AP latency (ms) | 108.4 | 43.3 | 37.7 | 14.9 | Paired t test | <0.001 | * |
| AP threshold (mV) | -38.4 | 3.7 | -41.5 | 4.8 | Paired t test | <0.001 | * |
| AP amplitude (mV) | 110.4 | 9.4 | 103.0 | 11.9 | Paired t test | <0.001 | * |
| FWHM (ms) | 1.32 | 0.22 | 2.37 | 0.6 | Wilcoxon test | <0.001 | * |
| Rise time (ms) | 0.29 | 0.04 | 0.35 | 0.1 | Wilcoxon test | <0.001 | * |
| Decay time (ms) | 0.68 | 0.12 | 1.35 | 0.4 | Wilcoxon test | <0.001 | * |

| | WT | SD | Fmr1KO | SD | Statistical test | P-value | |
|---|---|---|---|---|---|---|---|
| n | 26 | | 26 | | | | |
| RMP (mV) | -78.3 | 6.3 | -82.0 | 6.8 | Welch's t test | 0.0457 | * |
| Rin (MΩ) | 172.7 | 34.7 | 124.5 | 33.2 | Welch's t test | <0.001 | * |
| Tau (ms) | 14.8 | 4.8 | 12.0 | 4.2 | Mann-Whitney test | 0.0307 | * |
| Rheobase (pA) | 187.2 | 39.8 | 262.9 | 84.6 | Welch's t test | <0.001 | * |
| AP latency (ms) | 78.4 | 24.4 | 108.4 | 43.3 | Welch's t test | 0.0037 | * |
| AP threshold (mV) | -36.1 | 2.4 | -38.4 | 3.7 | Welch's t test | 0.0099 | * |
| AP amplitude (mV) | 103.5 | 11.7 | 110.4 | 9.4 | Unpaired t test | 0.022 | * |
| FWHM (ms) | 1.41 | 0.21 | 1.32 | 0.22 | Unpaired t test | 0.151 | |
| Rise time (ms) | 0.31 | 0.03 | 0.29 | 0.04 | Unpaired t test | 0.0246 | * |
| Decay time (ms) | 0.72 | 0.12 | 0.68 | 0.12 | Welch's t test | 0.2461 | |

| | WT | SD | Fmr1KO $BaCl_2$ | SD | Statistical test | P-value | |
|---|---|---|---|---|---|---|---|
| n | 26 | | 26 | | | | |
| RMP (mV) | -78.3 | 6.3 | -77.2 | 8.9 | Welch's t test | 0.87 | |
| Rin (MΩ) | 172.7 | 34.7 | 191.1 | 65.0 | Welch's t test | 0.0383 | * |
| Tau (ms) | 14.8 | 4.8 | 34.1 | 14.4 | Mann-Whitney test | 0.563 | |
| Rheobase (pA) | 187.2 | 39.8 | 178.8 | 81.1 | Mann-Whitney test | 0.163 | |
| AP latency (ms) | 78.4 | 24.4 | 37.7 | 14.9 | Welch's t test | <0.001 | |
| AP threshold (mV) | -36.1 | 2.4 | -41.5 | 4.8 | Welch's t test | 0.0138 | * |
| AP amplitude (mV) | 103.5 | 11.7 | 103.0 | 11.9 | Welch's t test | 0.561 | |
| FWHM (ms) | 1.41 | 0.21 | 2.37 | 0.6 | Mann-Whitney test | <0.001 | |
| Rise time (ms) | 0.31 | 0.03 | 0.35 | 0.1 | Mann-Whitney test | 0.0979 | |
| Decay time (ms) | 0.72 | 0.12 | 1.35 | 0.4 | Mann-Whitney test | <0.001 | |

Table 2. EPSP parameter ($BaCl_2$)

| | WT | SD | WT $BaCl_2$ | SD | Statistical test | P-value |
|---|---|---|---|---|---|---|
| n | | | | 25 | | |
| Frequency (Hz) | 3.56 | 0.89 | 3.65 | 0.91 | Paired t test | 0.278 |
| Amplitude (mV) | 0.68 | 0.12 | 0.72 | 0.16 | Paired t test | 0.0755 |
| n | | | | 24 | | |
| Sum ratio | 1.24 | 0.22 | 1.17 | 0.18 | Paired t test | 0.0921 |

| | Fmr1KO | SD | Fmr1KO $BaCl_2$ | SD | Statistical test | P-value |
|---|---|---|---|---|---|---|
| n | | | | 25 | | |
| Frequency (Hz) | 4.45 | 1.11 | 4.41 | 1.06 | Paired t test | 0.656 |
| Amplitude (mV) | 0.63 | 0.14 | 0.68 | 0.20 | Paired t test | 0.0425 |
| n | | | | 24 | | |
| Sum ratio | 1.00 | 0.26 | 1.08 | 0.23 | Paired t test | 0.259 |

| | WT | SD | Fmr1KO | SD | Statistical test | P-value | |
|---|---|---|---|---|---|---|---|
| n | | | 25 | | | | |
| Frequency (Hz) | 3.56 | 0.89 | 4.45 | 1.11 | Welch's t test | 0.003 | * |
| Amplitude (mV) | 0.68 | 0.12 | 0.63 | 0.14 | Welch's t test | 0.126 | |
| n | | | | 24 | | | |
| Sum ratio | 1.24 | 0.22 | 1.00 | 0.26 | Welch's t test | <0.001 | * |

| | WT | SD | Fmr1KO $BaCl_2$ | SD | Statistical test | P-value | |
|---|---|---|---|---|---|---|---|
| n | | | 25 | | | | |
| Frequency (Hz) | 3.56 | 0.89 | 4.41 | 1.06 | Welch's t test | 0.0037 | * |
| Amplitude (mV) | 0.68 | 0.11 | 0.67 | 0.19 | Welch's t test | 0.892 | |
| n | | | | 24 | | | |
| Sum ratio | 1.24 | 0.22 | 1.08 | 0.23 | Mann-Whitney test | 0.0105 | * |

Table 3. $Ba^{2+}$ sensitive current parameters

| | | | | | | | |
|---|---|---|---|---|---|---|---|
| | WT | SD | WT $BaCl_2$ | SD | Statistical test | P-value | |
| n | 8 | | 8 | | | | |
| Imax at -130 (pA) | -365.9 | 82.5 | -147.7 | 48.2 | Paired t test | <0.001 | * |
| | Fmr1KO | SD | Fmr1KO $BaCl_2$ | SD | Statistical test | P-value | |
| n | 10 | | 10 | | | | |
| Imax at -130 (pA) | -399.9 | 106.6 | -92.5 | 43.8 | Paired t test | <0.001 | * |
| | WT | SD | Fmr1KO | SD | Statistical test | P-value | |
| n | 8 | | 10 | | | | |
| Imax at -130 (pA) | -407.1 | 50.6 | -411.7 | 119.4 | Welch's t test | 0.59 | |
| | WT | SD | Fmr1KO $BaCl_2$ | SD | Statistical test | P-value | |
| n | 8 | | 10 | | | | |
| Imax at -130 (pA) | -407.1 | 50.6 | -94.1 | 46.6 | Welch's t test | <0.001 | * |
| | WT-WT $BaCl_2$ | | Fmr1KO - Fmr1KO BaCl | | | | |
| n | 8 | | 10 | | | | |
| I, % of inhibition | 59.6 | 12.0 | 76.9 | 7.6 | Welch's t test | 0.0045 | * |
| | WT | | Fmr1-KO | | | | |
| n | 8 | | 10 | | | | |
| Subtracted current (pA | -218.2 | 70.2 | -307.5 | 89.2 | Welch's t test | 0.0302 | * |
| Erev (mV) | -89.6 | 2.4 | -82.9 | 16.3 | Welch's t test | -0.126 | |

Table 4. Intrinsic excitability (ZD-7288)

| | WT | SD | WT ZD7288 | SD | Statistical test | P-value | |
|---|---|---|---|---|---|---|---|
| n | | | 17 | | | | |
| RMP (mV) | -78.4 | 7.4 | -83.1 | 6.3 | Paired t test | <0.001 | * |
| Rin (MΩ) | 132.0 | 34.5 | 140.0 | 31.0 | Paired t test | 0.0349 | * |
| Tau (ms) | 10.8 | 4.7 | 10.3 | 2.8 | Paired t test | 0.516 | |
| Rheobase (pA) | 226.6 | 76.2 | 256.4 | 86.3 | Paired t test | 0.0409 | * |
| AP latency (ms) | 94.2 | 90.7 | 87.1 | 35.2 | Paired t test | 0.719 | |
| AP threshold (mV) | -38.3 | 4.1 | -37.6 | 4.6 | Paired t test | 0.54 | |
| AP amplitude (mV | 102.6 | 14.1 | 105.0 | 14.6 | Paired t test | 0.148 | * |
| FWHM (ms) | 1.41 | 0.25 | 1.55 | 0.29 | Paired t test | <0.001 | * |
| Rise time (ms) | 0.37 | 0.09 | 0.41 | 0.12 | Paired t test | 0.0343 | * |
| Decay time (ms) | 0.72 | 0.14 | 0.79 | 0.15 | Paired t test | 0.002 | * |
| Sag (mV) | 1.83 | 1.37 | 1.04 | 0.34 | Paired t test | 0.0257 | * |

| | Fmr1KO | SD | Fmr1KO ZD728 | SD | Statistical test | P-value | |
|---|---|---|---|---|---|---|---|
| n | | | 17 | | | | |
| RMP (mV) | -84.1 | 6.7 | -87.4 | 6.3 | Paired t test | <0.001 | * |
| Rin (MΩ) | 92.3 | 36.1 | 104.4 | 36.5 | Paired t test | <0.001 | * |
| Tau (ms) | 6.4 | 1.5 | 7.5 | 2.1 | Paired t test | 0.0063 | * |
| Rheobase (pA) | 318.1 | 80.3 | 280.6 | 68.7 | Paired t test | <0.001 | * |
| AP latency (ms) | 144.3 | 95.3 | 92.4 | 44.7 | Paired t test | 0.0215 | * |
| AP threshold (mV) | -38.3 | 4.1 | -37.7 | 4.6 | Paired t test | 0.945 | |
| AP amplitude (mV | 114.4 | 10.6 | 115.7 | 13.2 | Paired t test | 0.543 | |
| FWHM (ms) | 1.23 | 0.14 | 1.34 | 0.16 | Paired t test | 0.0042 | * |
| Rise time (ms) | 0.28 | 0.03 | 0.30 | 0.04 | Paired t test | 0.0924 | |
| Decay time (ms) | 0.63 | 0.08 | 0.68 | 0.08 | Paired t test | 0.0128 | * |
| Sag (mV) | 1.53 | 0.49 | 1.14 | 0.61 | Paired t test | 0.0061 | * |

| | WT | SD | Fmr1KO | SD | Statistical test | P-value | |
|---|---|---|---|---|---|---|---|
| n | | | 17 | | | | |
| RMP (mV) | -78.4 | 7.5 | -84.1 | 6.7 | Unpaired t test | 0.0238 | * |
| Rin (MΩ) | 132.0 | 34.5 | 92.3 | 36.1 | Unpaired t test | 0.0025 | * |
| Tau (ms) | 10.8 | 4.7 | 6.4 | 1.5 | Welch's t test | 0.0014 | * |
| Rheobase (pA) | 226.6 | 76.2 | 318.1 | 80.3 | Welch's t test | 0.0018 | * |
| AP latency (ms) | 94.2 | 90.7 | 144.3 | 95.3 | Welch's t test | 0.126 | |
| AP threshold (mV) | -38.3 | 4.1 | -39.3 | 2.5 | Unpaired t test | 0.379 | |
| AP amplitude (mV | 102.6 | 14.1 | 114.4 | 10.6 | Welch's t test | 0.0097 | * |
| FWHM (ms) | 1.41 | 0.25 | 1.23 | 0.14 | Welch's t test | 0.0162 | |
| Rise time (ms) | 0.37 | 0.09 | 0.28 | 0.03 | Unpaired t test | <0.001 | * |
| Decay time (ms) | 0.72 | 0.14 | 0.63 | 0.08 | Welch's t test | 0.0265 | * |
| Sag (mV) | 1.83 | 1.37 | 1.53 | 0.49 | Welch's t test | 0.407 | |

| | WT | SD | Fmr1KO ZD728 | Sd | Statistical test | P-value | |
|---|---|---|---|---|---|---|---|
| n | | | | 17 | | | |
| RMP (mV) | -78.4 | 7.5 | -87.4 | 6.3 | Welch's t test | <0.001 | * |
| Rin (MΩ) | 132.0 | 34.5 | 104.4 | 36.5 | Unpaired t test | 0.0308 | * |
| Tau (ms) | 10.8 | 4.7 | 7.5 | 2.1 | Welch's t test | 0.0124 | * |
| Rheobase (pA) | 226.6 | 76.2 | 280.6 | 68.7 | Welch's t test | 0.0376 | * |

Table 4. Intrinsic excitability (ZD-7288) Continuation

| | | | | | | | |
|---|---|---|---|---|---|---|---|
| AP latency (ms) | 94.2 | 90.7 | 92.4 | 44.7 | Mann-Whitney test | 0.159 | |
| AP threshold (mV) | -38.3 | 4.1 | -39.3 | 2.5 | Unpaired t test | 0.385 | |
| AP amplitude (mV) | 102.6 | 14.1 | 115.7 | 13.2 | Welch's t test | 0.0087 | * |
| FWHM (ms) | 1.41 | 0.25 | 1.34 | 0.16 | Welch's t test | 0.361 | |
| Rise time (ms) | 0.37 | 0.09 | 0.30 | 0.04 | Mann-Whitney test | 0.0048 | * |
| Decay time (ms) | 0.72 | 0.14 | 0.68 | 0.08 | Welch's t test | 0.305 | |
| Sag (mV) | 1.83 | 1.37 | 1.14 | 0.61 | Welch's t test | 0.0703 | |

Table 5. EPSP parameters (ZD-7288)

| | WT | SD | WT ZD7288 | SD | Statistical test | P-value | |
|---|---|---|---|---|---|---|---|
| n | | | | 17 | | | |
| Frequency (Hz) | 2.45 | 0.94 | 2.73 | 1.12 | Paired t test | 0.0029 | * |
| Amplitude (mV) | 0.68 | 0.12 | 0.69 | 0.09 | Paired t test | 0.594 | |
| n | | | | 14 | | | |
| Sum ratio | 1.17 | 0.28 | 1.26 | 0.29 | Paired t test | 0.268 | |

| | Fmr1KO | SD | Fmr1KO ZD7288 | SD | Statistical test | P-value | |
|---|---|---|---|---|---|---|---|
| n | | | | 17 | | | |
| Frequency (Hz) | 2.97 | 1.01 | 3.98 | 1.38 | Paired t test | 0.0015 | * |
| Amplitude (mV) | 0.71 | 0.13 | 0.76 | 0.17 | Wilcoxon test | 0.0797 | |
| n | | | | 14 | | | |
| Sum ratio | 1.13 | 0.18 | 1.41 | 0.43 | Paired t test | 0.0086 | * |

| | WT | SD | Fmr1KO | SD | Statistical test | P-value |
|---|---|---|---|---|---|---|
| n | | | | 17 | | |
| Frequency (Hz) | 2.46 | 0.95 | 2.98 | 1.01 | Welch's t test | 0.132 |
| Amplitude (mV) | 0.68 | 0.12 | 0.71 | 0.13 | Welch's t test | 0.4544 |
| n | | | | 14 | | |
| Sum ratio | 1.17 | 0.28 | 1.13 | 0.18 | Unpaired t test | 0.668 |

| | WT | SD | Fmr1KO ZD7288 | SD | Statistical test | P-value | |
|---|---|---|---|---|---|---|---|
| n | | | | 17 | | | |
| Frequency (Hz) | 2.46 | 0.95 | 3.98 | 1.39 | Welch's t test | <0.001 | * |
| Amplitude (mV) | 0.68 | 0.12 | 0.76 | 0.17 | Welch's t test | 0.1259 | |
| n | | | | 14 | | | |
| Sum ratio | 1.17 | 0.28 | 1.41 | 0.43 | Mann-Whitney test | 0.0497 | * |
